# Astrocytes promote a protective immune response to brain *Toxoplasma gondii* infection via IL-33-ST2 signaling

**DOI:** 10.1101/338400

**Authors:** Katherine M. Still, Samantha J. Batista, Carleigh O’Brien, Oyebola O. Oyesola, Simon P. Früh, Lauren M. Webb, Igor Smirnov, Michael A. Kovacs, Maureen N. Cowan, Nikolas W. Hayes, Jeremy A. Thompson, Elia D. Tait-Wojno, Tajie H. Harris

## Abstract

Understanding how invading pathogens are sensed within the brain is necessary to uncover how effective immune response are mounted in immunoprivileged sites. The eukaryotic parasite *Toxoplasma gondii* colonizes the brain of its hosts and initiates robust immune cell recruitment, but little is known about innate recognition of *T. gondii* within brain tissue. The host damage signal IL-33 is one protein that has been implicated in control of chronic *T. gondii* infection, but the specific impact of IL-33 signaling within the brain is unclear. Here, we show that IL-33 is expressed by oligodendrocytes and astrocytes during *T. gondii* infection, is released into the cerebrospinal fluid of *T. gondii*-infected animals, and is required for control of infection. IL-33 signaling promotes chemokine expression within brain tissue and is required for the recruitment of peripheral anti-parasitic immune cells, including IFN-γ-expressing T cells and iNOS-expressing monocytes. Importantly, we find that the beneficial effects of IL-33 during chronic infection are not a result of signaling on infiltrating immune cells, but rather on radio-resistant responders, and specifically, astrocytes. Mice with IL-33R-deficient astrocytes fail to promote an adaptive immune response in the CNS and control parasite burden, demonstrating that astrocytes can directly respond to IL-33 *in vivo*. Together, these results indicate a brain-specific mechanism by which IL-33 is released and sensed locally, to engage the peripheral immune system in controlling a neurotropic pathogen.

## INTRODUCTION

Recruitment of immune cells to the brain during infection is a highly orchestrated process, requiring concerted expression of a number of chemokines and adhesion factors at the blood-brain barrier^1^. But the cues which precede these factors are less well understood. In particular, in many cases, it is unclear if brain resident cells possess the machinery to detect the presence of pathogens to promote the recruitment of peripheral cells. Murine infection with the eukaryotic parasite *Toxoplasma gondii (T. gondii)* features continual recruitment of blood-derived immune cells to the brain and serves as an excellent model for better understanding immune responses at this site.

*T. gondii* is a globally relevant pathogen which infects most warm-blooded vertebrates, including one-third of the human population^2-4^. Upon initial infection, *T. gondii* disseminates throughout peripheral tissues, where it is largely cleared, but ultimately persists in the brain of its hosts for their lifetime^2,5-8^. Mortality from *T. gondii* infection is associated with an increased prevalence of replicating parasite in brain tissue, documented in immunosuppressed patients undergoing transplant surgeries^9^, and in HIV-AIDS patients^10-12^, highlighting the importance of the immune response in controlling *T. gondii*. Indeed, control of brain *T. gondii* infection requires a Th1-dominated immune response,^2,8^ whereby CD4+ and CD8+ T cells and the IFN-γ they produce are required for survival^13^. Macrophages also exhibit anti-parasitic effector mechanisms which are necessary to control the parasite^8,14-20^.

It is not known, however, how the parasite is sensed in the brain to create an environment that promotes immune cell entry, stimulation, and maintenance. During the acute phase of infection in the periphery, dendritic cells and macrophages can sense either the parasite itself or host signals to initiate chemokine and cytokine expression which recruits and skews a strong Th1 immune response^8,21-25^. However, resident dendritic cells and peripheral immune cells do not exist in brain tissue under steady-state conditions^26,27^ and it is unclear if *T.gondii*-specific molecular patterns are sensed in this tissue. During chronic *T. gondii* infection of the brain, necrotic lesions form, characterized by the presence of replicating parasite, loss of brain-resident cell markers, and infiltration of immune cells, suggestive of tissue damage and alarmin release^13,28-32^. We hypothesized that indirect sensing of *T. gondii* infection, via recognition of host cell damage caused by the parasite, is an important step in instructing the immune response to *T. gondii* in the brain. Here we focused on the nuclear alarmin, IL-33, as a candidate orchestrator of the immune response to *T. gondii*. IL-33 is highly expressed in brain tissue^33^, and IL-33 signaling has been shown to be protective against tissue pathology during chronic *T. gondii* infection, but the mechanism by which IL-33 signals, and the immune mechanisms underlying this protection, were not studied in detail^34^.

IL-33 is categorized as an alarmin because it is known to amplify immune responses upon signaling through its receptor ST2, without a requirement for secretion or cleavage. A role for IL-33 in stroke^35,36^, neurodegeneration^37,38^, EAE^39,40^, and CNS infection^41-43^ has been described, but mechanistic understanding of IL-33 signaling during brain disease is limited. It is unclear in many instances which cell type(s) is the relevant responder to IL-33 during disease in the brain. While brain-resident macrophages have been shown to express IL-33 receptor at baseline^44^, astrocytes have been shown to upregulate it during pathology^36,45^. In addition, IL-33 receptor is also expressed on immune cells which can infiltrate the brain from the blood during disease^46,47^. Here, we separate the relative contribution of astrocytes, microglia/macrophages, and infiltrating immune cells in response to IL-33 release during brain pathology, finding that IL-33 can signal strictly within the brain to promote protective immunity to *T. gondii*.

## RESULTS

### IL-33 is released during *T. gondii* brain infection and is required to control parasite

To study *T. gondii* brain infection, we infected mice by intra-peritoneal injection with the avirulent *T. gondii* strain Me49 and waited four weeks post infection for a natural, chronic brain infection to be established. At this timepoint, peripheral infection has been controlled^6,48^. For all experiments, brain tissue was harvested at four weeks post infection unless otherwise specified.

Consistent with its role as a pre-stored alarmin, IL-33 is expressed highly at baseline^33,44,45^, and we detected only mild increases in IL-33 levels in infected versus uninfected tissue (Figure S1A). In the *T. gondii* infected brain parenchyma, we found nuclear IL-33 to be expressed by mature, CC1+ oligodendrocytes and by astrocytes, the percentages of which varied by brain region (Figures 1A, 1B, and S1B). While IL-33 was expressed almost exclusively by oligodendrocytes in white matter tracts, such as the corpus callosum, IL-33 expression in the cortex was split more evenly between astrocytes and oligodendrocytes (Figures 1A and 1B). Collectively, these results indicate that IL-33 is expressed by glia in the *T. gondii* infected mouse brain parenchyma. Importantly, we also detected IL-33 protein expression in astrocytes in healthy human brain tissue, as has been shown previously on the transcript level^49^(Figures S1C and S1D).

**Figure 1.**
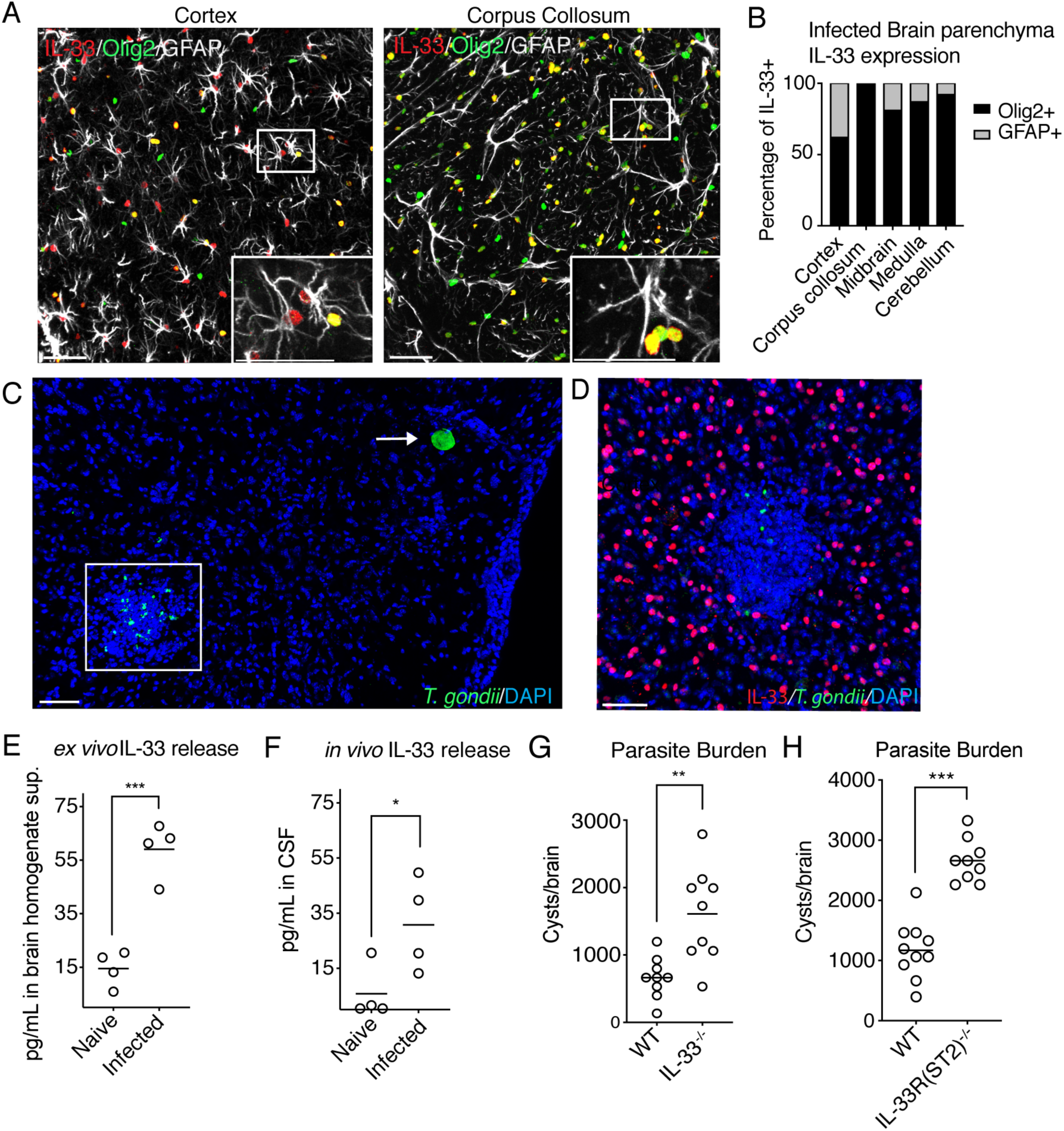
IL-33 is expressed by oligodendrocytes and astrocytes during brain *T. gondii* infection, is released during infection, and is required for control of parasite. **(A and B)** Representatitive images (A) and percentage quantification (B) of nuclear IL-33 expression (red) by cell type four weeks post-infection in various brain regions by confocal fluorescent microscopy. IL-33 is costained with the nuclear oligodendrocyte marker, Olig2 (green) or activated astrocyte marker, GFAP, (white). **(C and D)** Representative images of *T. gondii* in brain tissue, either in cyst form (arrow), or reactivated individual parasites (box)(C). IL-33-positive cells are absent from inflammatory foci containing replicating parasites (D). **(E and F)** Extracellular IL-33 release as measured by ELISA after *ex vivo* incubation of all cells isolated from infected brain tissue (E), or from CSF samples, each dot represents pooled CSF from 4-5 individual mice from a separate infection, displaying four infections in total (F). **(G and H)** Parasite burden as measured by cyst count from brain homogenate of IL-33-deficient (G) and IL-33 receptor (ST2)-deficient mice (H). Statistical significance was determined by two-tailed t-test (E) or randomized block ANOVA, when results from multiple independent experiments are shown (F-H). *= p<.05, **= p<.01, ***= p<.001. Scale bars indicate 50μm.

Since IL-33 does not need to be cleaved to be active, it is classically thought of as an alarmin that is released by necrotic cell death. During *T. gondii* infection, the parasite itself as well as the inflammatory environment provide opportunity for host cell damage and alarmin release. *T. gondii* predominantly exists in the brain as an intracellular cyst form which is slow growing^7,48^ (Figure 1C) and does not appear to pose an immediate risk to cells, due to a lack of observed tissue destruction and inflammation surrounding cysts (Figure 1C). But, for reasons not fully understood, cysts can reactivate anywhere in the brain, releasing individual parasites previously contained within the cyst wall^7,29^ (Figure 1C). Individual parasites can invade surrounding cells, replicate, and either lyse the cell or form a new cyst^50^. The presence of individual replicating parasites is correlated with morbidity in humans, and is thought to cause necrotic lesions in immunocompromised patients^9-12^. Interestingly, we found clusters of immune cells, including T cells and macrophages, surrounding individual replicating parasites but not *T. gondii* cysts (Figures S2A, S2B, 1C). Therefore, we hypothesized that either lytic *T. gondii* replication^50^ or local inflammation can cause release of local damage signals, such as IL-33. Corroborating this hypothesis, while IL-33 tiles evenly throughout uninfected brain tissue^45^, we noted focal loss of IL-33 staining, as well as oligodendrocyte and astrocyte markers, at sites of parasite replication and release (Figures 1D, S2C-E).

In order to determine if IL-33 is released during brain *T. gondii* infection, we first utilized an *ex-vivo* assay to measure extracellular IL-33 during infection. We processed uninfected (naïve) and infected mouse brains down to a single cell suspension containing brain resident cells as well as immune cells and parasite. We then incubated the single cell suspensions for four hours at 37°C before taking the supernatant and measuring extracellular IL-33 by ELISA. Strikingly, IL-33 was present in detectable quantities from supernatants of infected brain samples, but not naïve controls, suggesting that *T. gondii* infection has the capacity to induce IL-33 release. We then validated IL-33 release *in vivo*, by sampling CSF from the cisterna magna of infected mice and found detectable IL-33 levels in pooled CSF samples (Figure 1F).

We next asked if IL-33 was required to control infection. We infected wildtype and IL-33^-/-^ mice and assessed their brains for parasite burden, enumerating a significantly increased number of cysts in IL-33-deficient mice (Figure 1G). We also detected an increased parasite burden in the brains of infected IL-33 receptor (ST2) – deficient mice (Figure 1H), demonstrating a role for extracellular IL-33 signaling during this infection. At day 10 post-infection, no parasite was detected in the peritoneal cavity of infected ST2^-/-^ or wildtype animals. Therefore, ST2^-/-^ mice are not delayed or defective in clearing parasite in the periphery. While neither IL-33^-/-^ or ST2^-/-^ mice succumbed to infection, these results demonstrate that extracellular IL-33 signaling plays a role in control of *T. gondii* brain infection.

### IL-33-ST2 signaling is required for adequate numbers of functional T cells during chronic *T. gondii* infection

In order to better understand how IL-33 signaling was protective during *T. gondii* infection, we focused on characterizing ST2^-/-^ mice from this point onward. We first profiled recruited immune cell populations in the brain, beginning with the adaptive immune response, because a strong, Th1-biased immune response is absolutely critical for control of chronic *T. gondii* infection^13^. In the absence of IL-33 signaling during infection, approximately one-third fewer total T cells were present in the brain by flow cytometry (Figures 2A and 2B), suggesting an inability to recruit or maintain these cells. Approximately 96% of T cells in *T. gondii* infected brains were positive for either CD4 or CD8 markers (Figure 2A), and there was no selective decrease in either of these subsets, but rather a general reduction in T cell counts in ST2^-/-^ mice (Figures 2A and 2B). In addition to a reduction in T cells, T cell function was impaired in the absence of IL-33 signaling. Fewer CD4+ and CD8+ T cells were proliferating in the brain (Figures 2C and 2D), and fewer of these cells were making the critical cytokine IFN-γ by *ex vivo* restimulation with PMA/ionomycin (Figures 2E and 2F). IFN-γ is critical because it induces widespread anti-parasitic changes, such as increasing chemokine production^51,52^, adhesion factor expression^53^, and intracellular killing mechanisms of infected cell types, such as macrophages^8^. We found that the chief sources of IFN-γ in the brain during our infection were CD4+ and CD8+ T cells, with minimal contribution from NK cells, an important source of the cytokine during early stages of acute infection^54^(Figure S3A). These results show that extracellular IL-33 signaling is required to maintain an adequate anti-parasitic T cell response in brain tissue. Importantly, T cell numbers and function were unaffected at baseline in spleens of ST2^-/-^ mice, in the spleen during chronic infection, as well as at sites of inflammation such as the peritoneal cavity during acute infection (Figure S3B-E). We did, however, note a decrease in T cell numbers in the blood during acute infection of ST2^-/-^ mice (Figure S3F).

**Figure 2.**
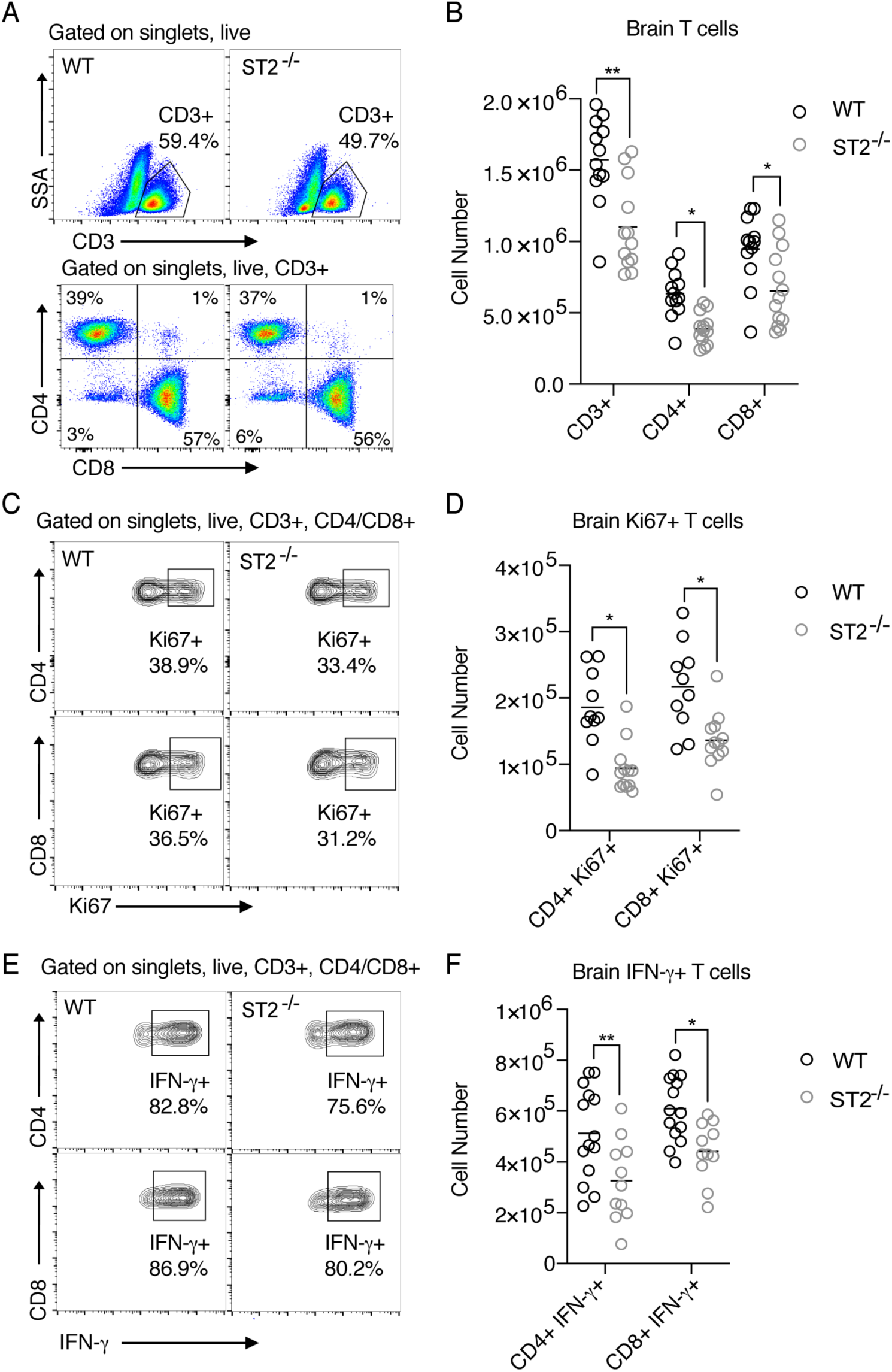
IL-33-ST2 signaling is required for adequate numbers and functionality of T cells in the brain during chronic *T. gondii* infection. **(A and B)** Representative flow cytometry plots (A) and quantification (B) of infiltrated total T cells in infected brain tissue four weeks-post infection. **(C and D)** Representative flow cytometry plots (C) and quantification (D) showing frequency of proliferating (Ki67+) T subsets in infected brain tissue. **(E and F)** Representative flow cytometry plots (E) and number (F) of IFN-γ-positive T cell subsets in infected brain tissue. IFN-γ was measured following *ex-vivo* re-stimulation of brain cells, incubated with Brefeldin A and PMA/ionomycin for 5 hours at 37°C (F). Statistical significance was determined by randomized block ANOVA (B,D,F), and each quantified panel displays three pooled independent experiments. *= p<.05, **= p<.01, ***= p<.001

### IL-33-ST2 signaling is required for the recruitment and anti-parasitic function of myeloid cells during brain *T. gondii* infection

We were also interested in the impact of IL-33 signaling on the myeloid cell lineage since macrophages cluster tightly around replicating parasite in brain tissue and express iNOS, a key molecule invovled in the control of *T. gondii* (Figure S2B)^8^. We assessed numbers of CD11b+ myeloid cells in infected ST2^-/-^ mice by flow cytometry, using CD45hi expression to differentiate infiltrating myeloid cells from CD45int microglia (Figures 3A and 3B). Many of the CD45hi CD11b+ cells recruited to the *T. gondii*-infected brain are Ly6C+ CD11c- and Ly6G-, indicating that a large portion of recruited myeloid cells are Ly6Chi monocytes and Ly6Clo monocyte-derived macrophages^55^. Importantly, infected ST2^-/-^ mice displayed a reduced frequency and number of CD45hi myeloid cells, by approximately half, in the brain during chronic infection, while CD45int cell numbers held constant (Figures 3A and 3B).

**Figure 3.**
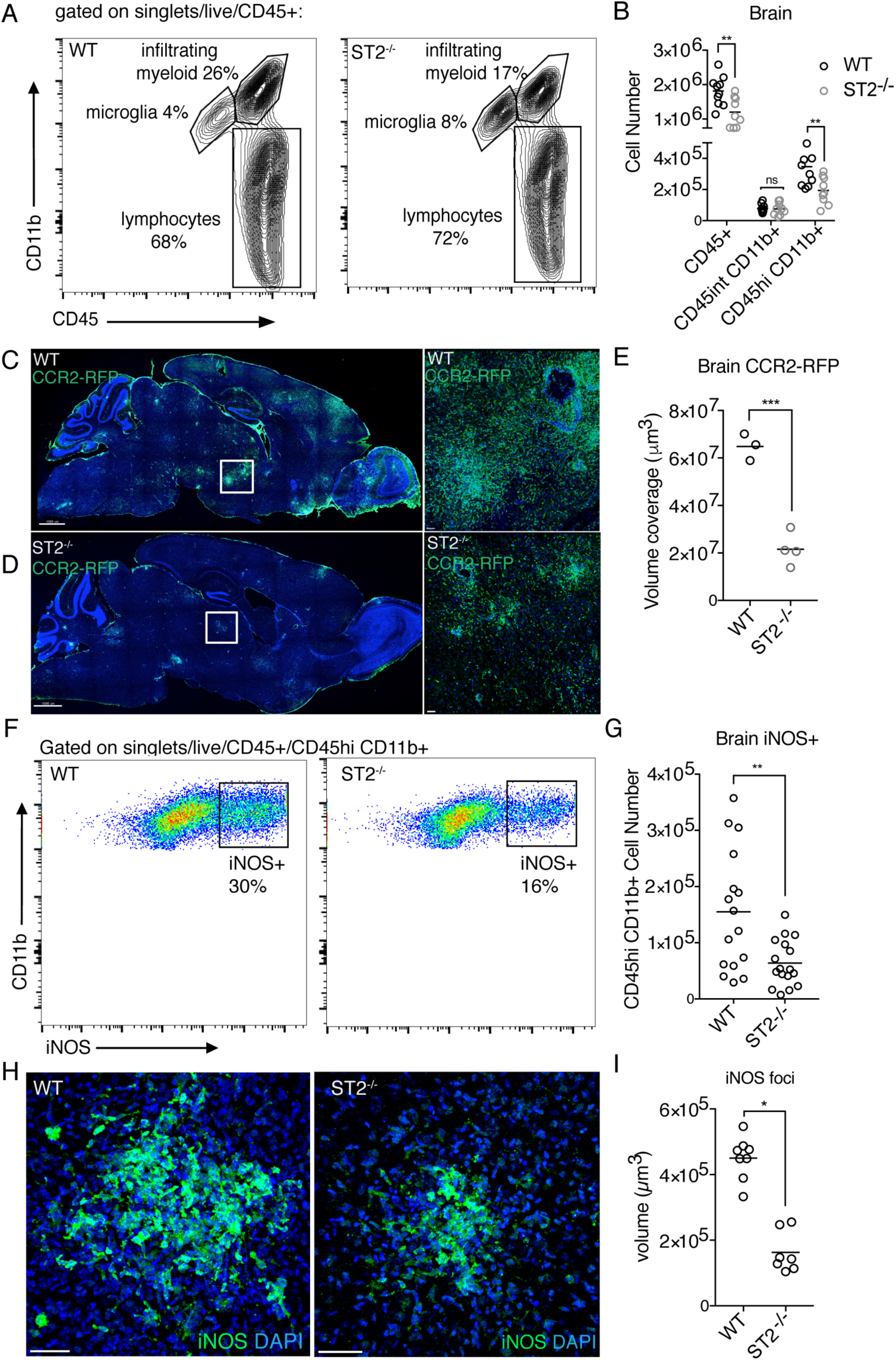
IL-33-ST2 signaling is required for the recruitment and anti-parasitic function of peripheral myeloid cells in the brain during chronic *T. gondi* infection. **(A and B)** Representative flow cytometry plots (A) and quantification (B) of CD11b+ myeloid cells in the brain 4 weeks post infection. CD45hi expression was used to differentiate infiltrating myeloid cells from CD45int microglia. **(C-E)** Visualization (C and D), and quantification (E) of infiltrated CCR2+ monocytes by fluorescence confocal microscopy of **i**nfected CCR2-RFP reporter mice. ST2-deficient mice were crossed to CCR2-RFP mice to assess the contribution of IL-33-ST2 signaling to monocyte recruitment. **(F and G)** Representative flow cytometry plots (F) and quantification (G) of iNOS+ CD45hi, CD11b+ infiltrating cells in the brain. **(H and I)** Visualization (H) and quantification (I) of the size of iNOS foci in brain tissue by fluorescence confocal microscopy. Statistical significance was determined by randomized block ANOVA when two or more experiments were pooled (B, G, I), and by a two-tailed t-test (E). *= p<.05, **= p<.01, ***= p<.001. Scale bars indicate 2000μm (C,D) and 50μm (H).

To more specifically assess monocyte recruitment, we used CCR2-RFP reporter mice to visualize monocytes in the brain during chronic infection. Monocyte-derived cells are critical for control of brain *T. gondii* infection, as anti-CCR2 antibody administered during chronic infection results in rapid mortality^55^. By immunohistochemistry, we observed large numbers of CCR2+ cells throughout the brain 4 weeks post infection in wildtype mice (Figure 3C). Importantly, we crossed CCR2-reporters to a ST2^-/-^ background, which revealed a marked reduction in CCR2+ cells (Figures 3D and 3E). These results confirmed that there is a defect in monocyte recruitment in the absence of IL-33 signaling during infection.

Next, we assessed the anti-parasitic function of the myeloid compartment in the absence of IL-33 signaling. We focused on inducible nitric oxide synthase (iNOS) expression. Synthesis of nitric oxide by host cells can deprive the parasite of essential amino acids and prevent parasite growth *in vitro*^56^. *In vivo*, iNOS knockout mice succumb to infection during the early chronic phase^15^. Of the myeloid cells that were able to infiltrate the brain, fewer of these cells were making iNOS in the absence of IL-33 signaling (Figures 3F-I). Interestingly, we did not detect significant iNOS positivity in any tissues, aside from the brain, in acute or chronic infection (Figure S4). Additionally, no defects in myeloid cell number were found at baseline in uninfected ST2^-/-^ mice, in peripheral tissues during acute infection, including in the peritoneum, blood, and spleen, or in peripheral tissues during chronic infection (Figure S4). In sum, these results, in conjunction with T cell deficits, demonstrate that IL-33 signaling impacts the presence and function of immune cell populations that are necessary for controlling *T. gondii* infection in the brain.

### IL-33-ST2 signaling induces expression of factors involved in recruitment of immune cells to the brain

Next, we sought to address how IL-33 is changing the environment within the brain to make the tissue conducive to immune cell recruitment. Initiation and maintenance of an immune response in the brain requires expression of cytokines, chemokines, adhesion factors, and factors which promote the entry of and maintain proliferation of immune cells^1^. It is well known that *T. gondii* infection induces many of these factors^51-53,57^,which we validated by whole brain qRT-PCR. We find a profound increase in brain *ccl2, cxcl9, cxcl10, ccl5, cxcl1, vcam*, and *icam* expression over uninfected controls (Figure 4A). When we assessed the same genes in infected ST2-deficient mice, we saw significantly decreased expression of the chemokines *ccl2, cxcl10*, and *cxcl1* (Figure 4A), along with smaller reductions in the expression of the adhesion factors *vcam* and *icam* (Figure 4B). Interestingly, *ccl2* and *cxcl10* expression has been attributed to astrocytes by *in situ* hybridization during *T. gondii* infection^52^. Although *cxcl1* has not been extensively studied in chronic *T. gondii* infection, it has also been reported in astrocytes during neuroinflammation^58^. However, we did not observe an effect of IL-33 signaling on *cxcl9* expression, a chemokine which is made by PU.1-expressing cells rather than astrocytes^59^ (Figure 4B). These results suggest that IL-33 signaling, either directly or indirectly, could be inducing chemokine expression in astrocytes during *T. gondii* infection.

**Figure 4.**
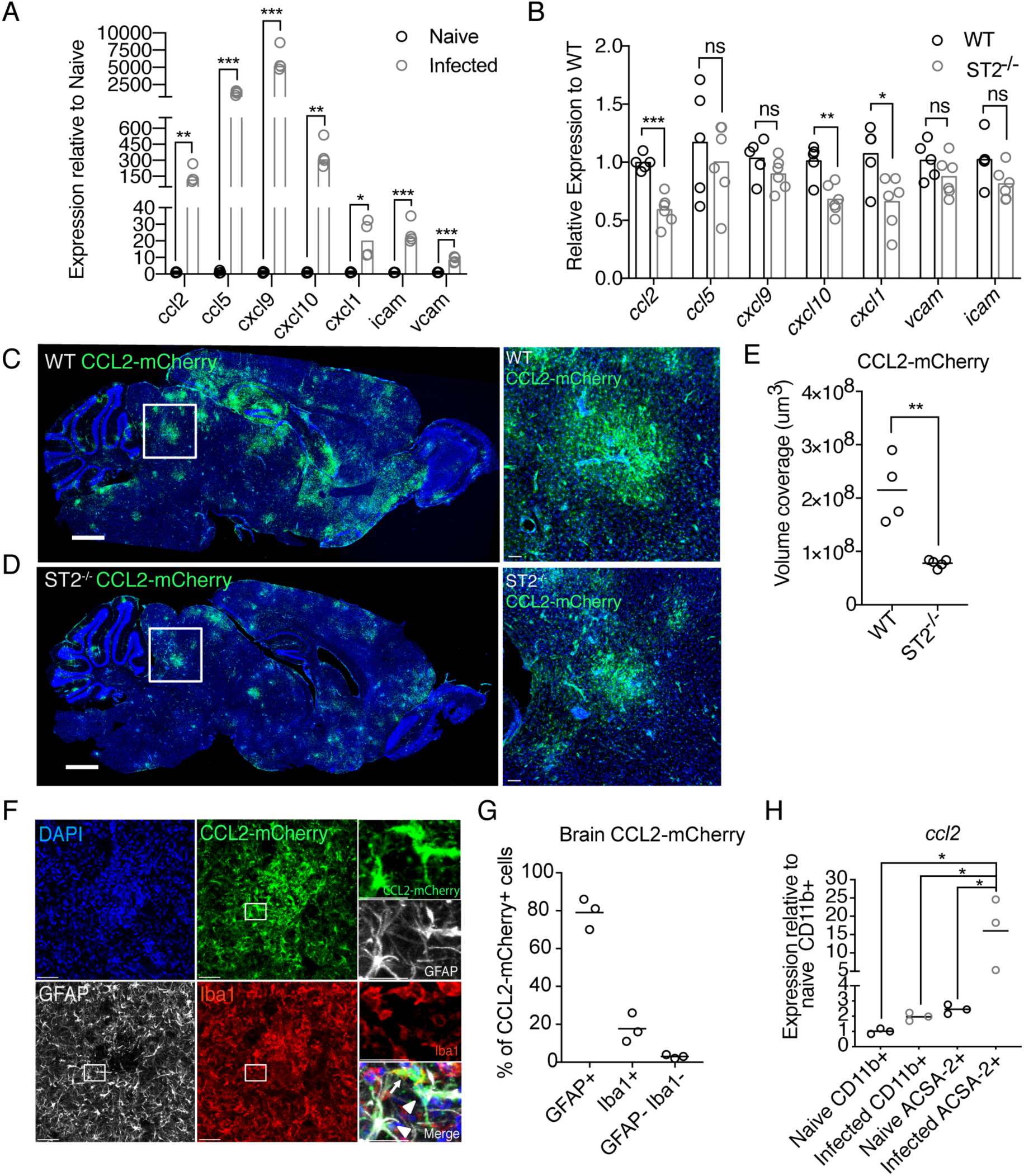
IL-33-ST2 signaling induces factors which recruit immune cells to the brain. **(A and B)** Real time PCR from whole brain homogenate was used to assess changes in chemokine and adhesion factor gene expression from naïve to infected animals (A) and between wildtype and ST2-deficient animals (B) at four weeks post infection. **(C-E)** Visualization (C,D) and quantification (E) of CCL2-RFP reporter expression by mCherry staining (green) in infected brain tissue by confocal fluorescence microscopy. **(F and G)** Breakdown of CCL2-mCherry reporter positivity by cell type using confocal microscopy. mCherry (green) is colocalized with Iba1+ macrophages (red) or activated, GFAP+ astrocytes (white). **(H)** Cell-type specific magnetic enrichment for myeloid cells (CD11b+) or astrocytes (CD11b- and ACSA-2+) in naïve and chronically infected brain tissue. Single cell suspensions of enriched cells were resuspended in Trizol, RNA extracted, and run by real time PCR for *ccl2* expression. Statistical significance was determined by two-tailed t-test (A, B, E) or One-way ANOVA with Tukey’s test (H). *= p<.05, **= p<.01, ***= p<.001. Scale bars indicate 2000μm (C,D) and 50μm (F).

We next wanted to identify the cellular source of chemokine in infected brain tissue to better understand where IL-33 was exerting its effects. We focused on studying the expression pattern of a chemokine whose transcript levels were altered to the greatest degree in the absence of IL-33 signaling, the monocyte chemoattractant *ccl2*. We used immunofluorescence microscopy to image the brain tissue of chronically infected *ccl2*-mCherry reporter mice^60^. We observed *ccl2-* mCherry expression in “hotspots” throughout the brain, implicating a local response to signals in brain tissue (Figure 4C). We validated by immunohistochemistry that *ccl2* expression was greatly reduced in ST2^-/-^ mice by crossing our *ccl2*-mCherry reporters to an ST2-deficient background (Figures 4C-4E). Specifically, *ccl2* foci in ST2^-/-^ mice were much reduced in size compared with wildtype infected mice (Figures 4D and 4E). We then assessed which cells were producing *ccl2. ccl2* in infected brain tissue colocalized with both GFAP+ astrocytes and Iba1+ macrophages (Figure 4F). Approximately 75% of cells expressing *ccl2* were astrocytes, 22% were Iba1+ macrophages, and 3% of cells did not co-stain with either of these markers (Figure 4G). These results were validated by assessing *ccl2* expression in brain cells magnetically enriched for either astrocytes or macrophages. To enrich for astrocytes, we used a previously validated protocol, which first removes CD11b+ cells by negative magnetic isolation, followed by positive isolation of ACSA-2 positive cells^61^. At the same time, we enriched for CD11b+ myeloid cells by keeping the CD11b+ magnetically-enriched fraction (Figure S5). RNA from these populations showed a 10-fold higher *ccl2* expression in astrocytes compared with macrophages during chronic infection (Figure 4H). Although multiple signals undoubtedly converge to induce chemokine expression such as *ccl2* during infection, including other innate cytokines^62^, our data suggest that IL-33 is a major contributor to the induction of local *ccl2*. These results support a role for IL-33 in inducing chemokine production to promote immune cell entry to the brain.

### IL-33 signals on a radio-resistant cell type to control chronic *T. gondii* infection

In order to further understand how IL-33 signaling supports protective immune responses, we began to narrow down the cell type that directly responds to IL-33 in the brain. In studies concerning IL-33 during CNS disease, genetic evidence for a responding cell type is lacking. This is likely due in part to the fact that several brain-resident cells can express ST2 during disease, such as microglia, astrocytes, and endothelial cells^36,63-65^, but a wide range of immune cells can also express the receptor, including but not limited to ILC2s, regulatory T cells, Th2 cells, mast cells, and macrophages^47,66,67^. We performed a bone marrow chimera to determine if IL-33 signals on a radio-sensitive or a radio-resistant cell type to exert its effects during chronic *T. gondii* infection. We lethally irradiated wildtype and ST2^-/-^ mice and reconstituted these mice with bone marrow from either wildtype donors or ST2^-/-^ donors (Figure 5A). Blood was assessed for reconstitution prior to infection (Figures 5B-D), and in all cases, immune cells were >90% positive for the congenic CD45 marker of donor mice, confirming successful reconstitution of the chimera (Figures 5B-D). We note that irradiation did alter the severity of infection, resulting in high parasite burdens in all groups, and a high number of infiltrating cells (Figures 5E-G). Nonetheless, these experiments revealed that ST2-deficiency on radio-resistant cells, recapitulated the phenotype of global knockouts, including reduced infiltrating myeloid and T cell numbers and parasite burden (Figures 5E-G).

**Figure 5.**
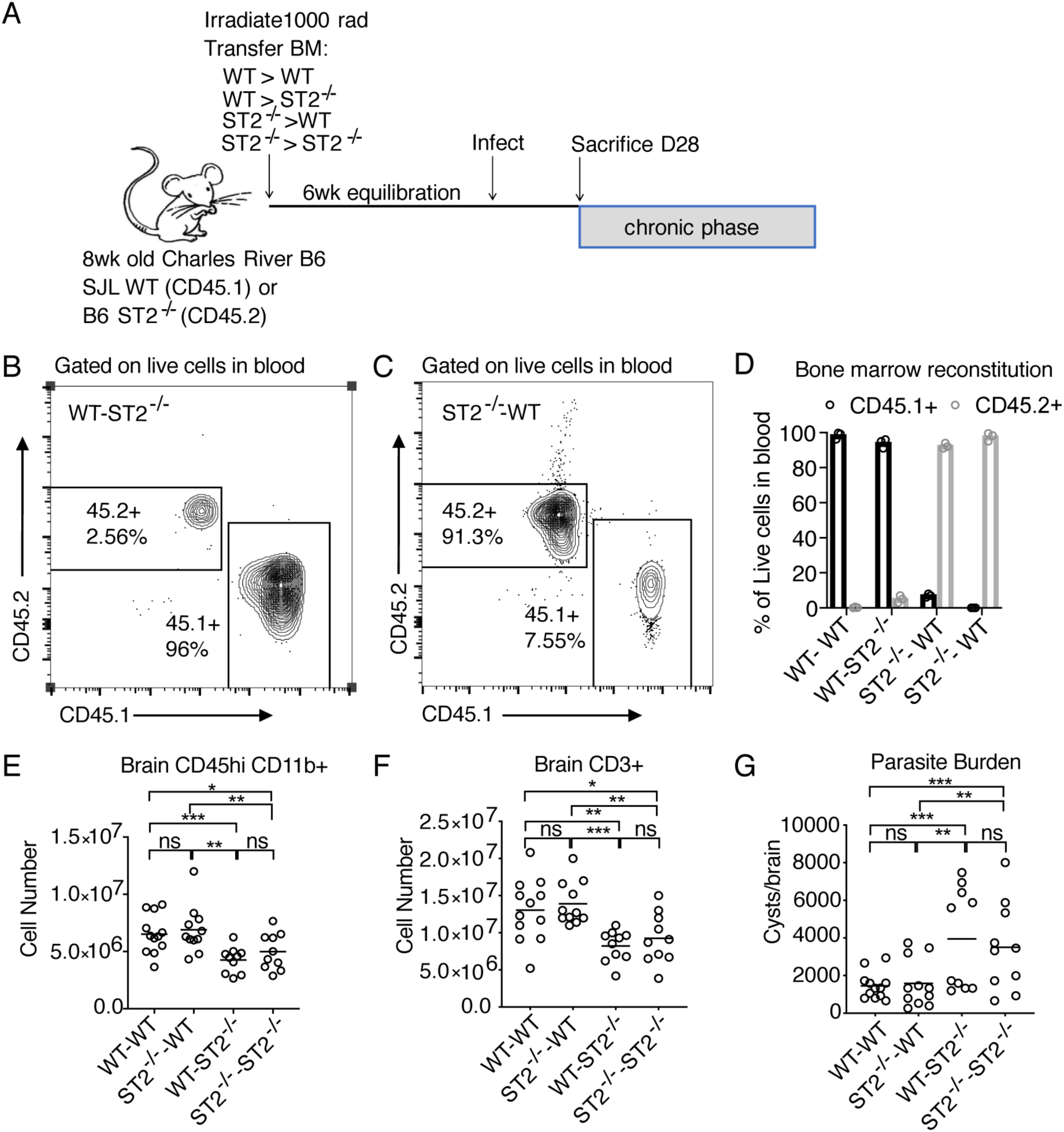
IL-33 signals on a radio-resistant responder. **(A)** Bone marrow chimera experimental setup. **(B-D)** Representative flow cytometry plots (B,C) and quantification (D) of bone marrow reconstitution in blood 6 weeks post irradiation and prior to infection. **(E and F)** Quantification of immune cells, including infiltrating myeloid cells (E) and T cells (F) in the brain at 4 weeks post infection following irradiation. **(G)** Parasite burden as assessed by cyst count of brain homogenate. Statistical significance was determined by randomized block ANOVA (E-G), each panel showing data pooled from two independent experiments *= p<.05, **= p<.01, ***= p<.001.

Importantly, microglia are not fully replenished by bone marrow-derived cells post-irradiation. Therefore, we cannot rule out microglia as responding to IL-33 in the brain. In addition, microglia highly express ST2 at baseline^36,44,45,63^, prior to chronic infection. To better understand which radio-resistant cells were capable of responding to IL-33, we magnetically enriched for CD11b+ cells (microglia/macrophages), and ACSA-2+ astrocytes from naïve and chronically infected brain tissue, and compared *il1rl1* (st2) expression. These cell types have been previously reported to express the IL-33 receptor, are major producers of IL-33-dependent chemokine during *T. gondii* infection, and are generally required for control of *T. gondii* infection. We found that microglia/macrophages cells express *il1rl1* at high levels in uninfected mice, but downregulate the IL-33 receptor 20-fold upon infection, while astrocytes express low levels at baseline and increase receptor expression with infection (Figure S6A), thus indicating a change in capability of cells which are able to respond to IL-33 prior to and following infection.

### IL-33-ST2 signaling on astrocytes, but not microglia/macrophages, potentiates immune responses and limits parasite burden

To determine if IL-33 directly signals on microglia/macrophages or astrocytes, we crossed ST2fl/fl mice^68^, to either constitutive CX3CR1cre mice or GFAPcre mice, respectively. We observed that deletion of ST2 from macrophages did not affect any major phenotype observed in ST2^-/-^ knockout mice, including brain myeloid cell number and function, T cell number, or parasite burden (Figure 6A-C). We observed minor defects in CD4+ T cell number in these mice, including CD4+ T cell proliferation (Figure 6B). Direct IL-33-ST2 signaling on astrocytes, however, did impact immune cell numbers in the brain, including infiltrating myeloid cell and T cells. T cell function, including proliferation and IFN-γ production, was also reduced in mice with IL-33R-deficient astrocytes (Figures 6D and 6E). Importantly, aligned with a decrease in T cell-derived IFN-γ, which is critical for control of infection^13^, GFAPcre ST2fl/fl mice displayed increased parasite burden (Figure 6F). Additionally, GFAPcre ST2fl/fl mice did not have defects in peripheral immune cell compartments, displaying increased immune cell numbers in the spleen during chronic infection (Figure S6B), to a level comparable with whole-body ST2-deficiency. Collectively, these results show that astrocytes respond to a damage signal, IL-33, to promote a protective immune response to *T. gondii* in the brain.

**Figure 6.**
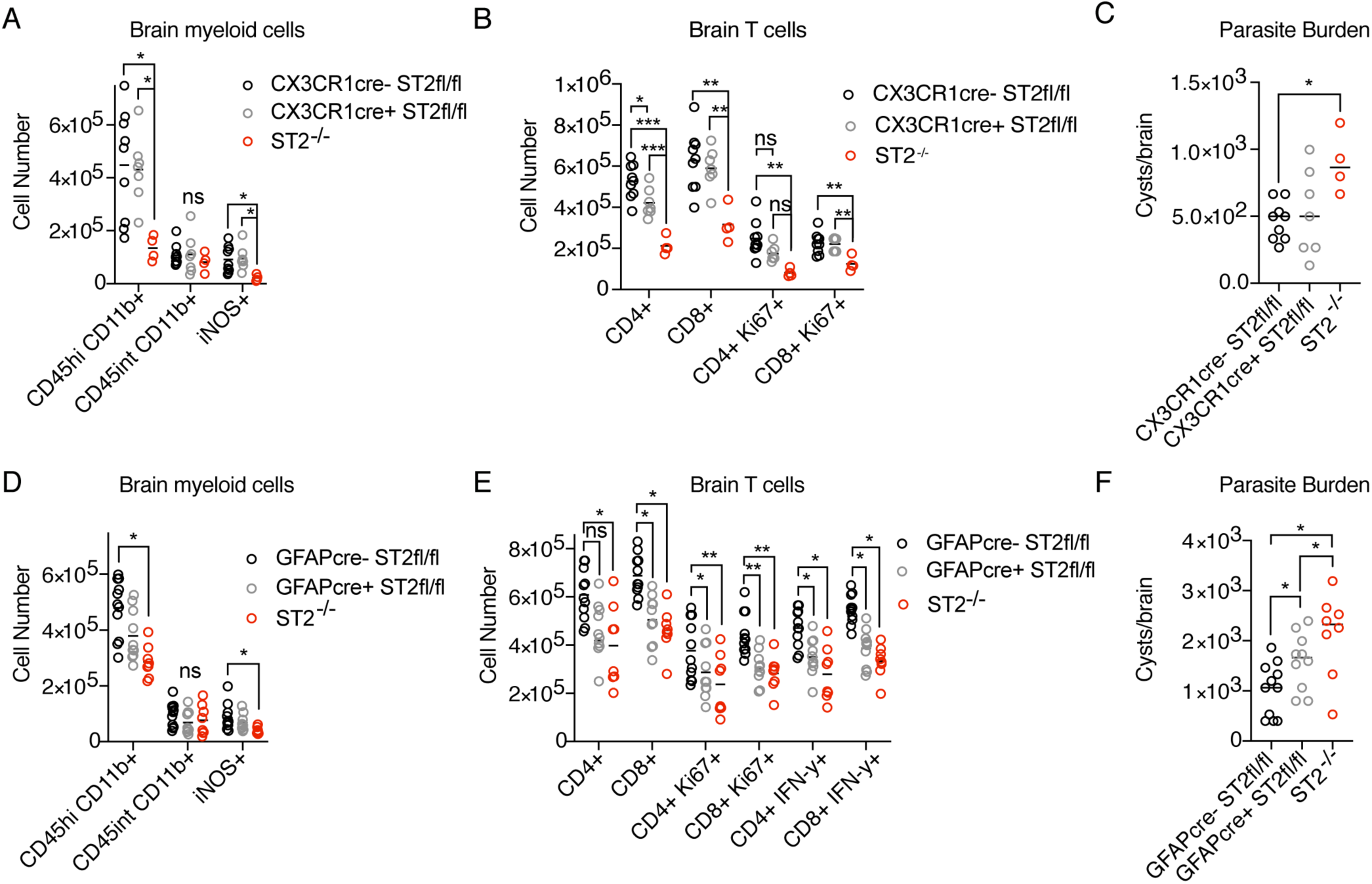
IL-33-ST2 signaling on astrocytes, but not macrophages, is required to control brain *T. gondii* infection. **(A and B)** Quantification of brain immune cells in CX3CR1cre ST2fl/fl mice by flow cytometry – including myeloid cells (A) and T cell subsets (B) at four weeks post infection. **(C)** Parasite burden in CX3CR1cre ST2fl/fl mice as assessed by cyst enumeration in brain homogenate. **(D and E)** Quantification of brain immune cells in GFAPcre ST2fl/fl mice by flow cytometry – including myeloid cells (D) and T cell subsets (E). IFN-γ was measured following *ex-vivo* re-stimulation of brain cells, incubated with Brefeldin A, and PMA/ionomycin for 5 hours at 37°C. **(F)** Parasite burden in GFAPcre ST2fl/fl mice as assessed by cyst enumeration in brain homogenate. Statistical significance was determined by one-way ANOVA with Tukey’s test (A-C), or by randomized block ANOVA(D-F), where each panel shows data pooled from two independent experiments *= p<.05, **= p<.01, ***= p<.001.

## DISCUSSION

We observe that brain resident cells can respond to IL-33, a damage-associated molecular pattern (DAMP), to initiate protective immunity during chronic *T. gondii* infection. This work begins to uncover the mechanisms by which the presence of *T. gondii* can be sensed in brain tissue, which is devoid of peripheral immune cells in the healthy state. Indeed, much of the work on sensing of *T. gondii* has focused on immune cell recognition of *T. gondii* in peripheral tissues, during the acute phase of infection. It has been shown that *T. gondii* profilin protein, a pathogen-associated molecular pattern (PAMP), can be sensed directly by TLR11 and TLR12-expressing dendritic cells in the periphery, which are capable of initiating protective Th1 immunity by producing IL-12^8,21,22^. In humans, however, TLR11 is a pseudogene and TLR12 is not expressed^69^. Thus, more recent studies have focused on innate sensing mechanisms with potential relevance to humans, including NOD-like receptor sensing of *T. gondii*^*23,24*^, as well as alarmin recognition, such as S100A11^25^, by macrophages. Since we find IL-33 to be highly expressed in human brain tissue, we hypothesize that our results will be relevant to control of human brain infection. Furthermore, our results shed light on the importance of continual DAMP recognition to contain infection long after adaptive immune responses are first primed in peripheral tissues.

We found nuclear IL-33 to be expressed by oligodendrocytes and astrocytes in the mouse *T. gondii*-infected brain, and we were able to detect extracellular IL-33 in the cerebrospinal fluid during chronic infection. Since extracellular IL-33 is negatively regulated in a number of ways to limit inflammation^70-72^, this result suggests high levels of IL-33 release during brain *T. gondii* infection. Our results also raise the question of how IL-33 is liberated from astrocytes and oligodendrocytes during disease, whether by cell death or secretion^67^. Mechanisms of glial death during disease have not been extensively studied *in vivo*, and it is unclear if the parasite itself, or if secondary, toxic inflammation can mediate the release of host damage signals. It is also unclear how long-ranging the effects of alarmin signaling are. Work on IL-33 in cardiac muscle^68^, as well as the retina^73^, has indicated that IL-33 often signals very locally, acting in either an autocrine or paracrine manner. For example, Müller cells, specialized astrocytes in the retina, have been implicated in the release and sensing of IL-33 in the retina^73^. Our work further corroborates local IL-33 signaling, as IL-33-dependent *ccl2* expression is found adjacent to necrotic lesions containing replicating *T. gondii*. Thus, IL-33 may be released by astrocytes or oligodendrocytes, and signal on the same cell type, including adjacent cells, to induce chemokine expression necessary for the recruitment of anti-parasitic immune cells to the brain.

In the lung, gut, and skin, IL-33 has historically been shown to signal almost exclusively on immune cells to orchestrate immunity^47^. However, recent work demonstrates that IL-33 can signal on non-hematopoeitic cells – including cardiomyocytes in the heart^68^ and endothelial cells *in vitro*. We found, consistent with a previous report in stroke^36^, that IL-33 signals on a radio-resistant responder during brain *T. gondii* infection. Tissue microenvironment and context likely influence whether IL-33 signaling on immune cells is relevant. Peripheral immune cells are not present in healthy brain tissue due to the blood brain barrier, thus necessitating the capability of brain resident cells to respond to damage. Additionally, the environmental milieu specific to CNS *T. gondii* infection promotes a robust Th-1skewed immune response necessary for intracellular parasite killing, characterized by T-cell derived IFN-γ and macrophage-derived iNOS^6,8^. Thus, the IL-33-dependent mechanisms, specifically those activating type 2 immune responses that drive asthma, allergy, and the expulsion of helminth infections^46,66^, may not be relevant to brain infection with an intracellular pathogen.

When we examined brain-resident cell expression of ST2, we found that microglia express high levels of ST2 in healthy brain tissue, but they downregulate the receptor 20-fold upon infection. A downshift in microglia inflammatory genes has been reported in other disease states, and it is unclear what factors drive this process. However, these results do not rule out the importance of IL-33 signaling on microglia in other settings. Indeed, during development, microglia expression of ST2 is required for microglial phagocytosis of synapses^44^. While we did not find any major impact of microglia/macrophage ST2 expression on control of chronic *T. gondii* infection, it is possible that microglia respond to IL-33 at earlier timepoints, such as when the parasite first reaches the brain. If microglia normally respond to IL-33 earlier in infection, our results indicate that astrocyte responses to IL-33, or additional MyD88-dependent signals, may compensate for the lack of ST2-signaling in microglia.

In contrast to microglia/macrophages, we found that astrocytes increase expression of the IL-33 receptor upon infection. Astrocytes are critical for controlling CNS infection and potentiate inflammation. For example, astrocyte responses to IFN-γ are necessary to control *T. gondii* infection^74^. We found that IL-33 receptor expression on astrocytes was required for adequate promotion of an anti-parasitic immune response in the brain, including proliferation and IFN-γ production in T cells, which is critical for control of *T. gondii*. This is the first time, to our knowledge, that astrocytes have been directly implicated in an IL-33 response via genetic manipulation. Our results also raise questions regarding the role of astrocytes in supporting T cell responses to infection and whether these responses are direct or indirect. IL-33 signals through the adaptor MyD88^66^, and could influence the expression of numerous molecules that support T cell responses, including trophic factors, TCR engagement, chemokines, or blood brain barrier opening for T cell entry.

Notably, immune defects in GFAPcre ST2fl/fl mice did not match the magnitude observed in ST2^-/-^ mice. These results either implicate compensation by other cell types when ST2 is deleted from a single population, or the presence of additional ST2-expressing radio-resistant cell types not considered here. One possibility is endothelial cells, which have recently been shown to express ST2 during disease and can be directly activated in response to IL-33 *in vitro*^*65,75*^. Nonetheless, our results indicate that astrocytes are capable of responding to extracellular IL-33 during infection. This finding, in addition to the detection of IL-33 release in the brain, has broad implications for sensing of tissue damage in other disease states, as definitive evidence that alarmins can signal strictly within brain tissue to initiate protective immunity.

Our results underscore the importance of the signaling of one alarmin in controlling chronic *T. gondii* infection. Little is known about the direct recognition of *T. gondii* in brain tissue and we implicate the response to infection-induced damage as a major mechanism of innate immune activation in the brain throughout chronic infection. Numerous canonical alarmins are expressed in the brain and their receptors are differentially expressed by cell type. Thus, there is likely a complex integration of multiple PAMP and DAMP signals that is required to allow peripheral immune cells to overcome the blood brain barrier, infiltrate the tissue parenchyma, and carry out effector function. Identifying the cell types that can repond to specific DAMPs in the brain will be key not only to understanding immunity to CNS infection but any neuroinflammatory condition with hallmarks of cell death, including neurodegeneration.

## METHODS

### Contact for reagent and resource sharing

Further information and requests for resources and reagents should be directed to and will be fulfilled by the Lead Contact, Tajie Harris.

### Experimental mice

C57BL/6, CCL2-RFP^flox^, CCR2^RFP^, GFAPcre, and CX3CR1cre mice were purchased from Jackson Laboratories. B6.SJL-Ptprc^a^ Pepc^b^/BoyCrCl (C57BL/6 Ly5.1) mice were purchased from Charles River. Ordering information for these strains is listed in the key resources table. ST2(il1rl1)^-/-^ mice were generously provided by Andrew McKenzie (Cambridge University). IL-33^-/-^ animals were obtained from Amgen by Elia Tait Wojno, currently at the University of Washington and previously at Cornell University. All IL-33-/- animals were housed at Cornell University. ST2(il1rl1)-floxed embryos were received from KOMP repository (RRID MMRRC:048182-UCD). Importantly, we received notice after fully breeding these mice to cre-expressing mice, that the deposited floxed embryos contain a copy of an inducible, cardiomyocyte-driven cre^68^, unknown to the KOMP repository. Upon receiving notice, we did detect the presence of MYH6cre^ERT2^ alleles throughout our ST2fl/fl colony. Importantly, we deleted ST2 under the control of constitutive cre drivers and never administered tamoxifen to these mice. We performed statistical tests to determine the effect of MYH6cre expression, and did not find evidence of an effect in experiments where the MYH6cre was present in some animals and not others. Moreover, in the experiments reported here (Figure 6), all animals were positive for the MYH6cre allele. All animals were housed in a UVA specific pathogen-free facility with a 12h light/dark cycle. Mice were age and sex matched for each experiment, and were sacrificed in parallel. Animals were infected with *T. gondii* at 7 to 9 weeks of age and were housed separately from breeding animals. All procedures adhered to regulations of the Institutional Animal Care and Use Committee (ACUC) at the University of Virginia and Cornell University.

### Human brain tissue

Healthy human brain samples from adult patients were obtained from the UVA Human Biorepository and Tissue Research Facility. Samples were preserved on paraffin embedded slides. Patient identification and medical background was withheld and therefore IRB approval was not required.

### Parasite strains

The avirulent, type II *Toxoplasma gondii* strain Me49 was used for all infections. *T. gondii* cysts were maintained in chronically infected (1-6 months) Swiss Webster (Charles River) mice. To generate cysts for experimental infections, CBA/J (Jackson Laboratories) mice were infected with 10 cysts from brain homogenate of Swiss Webster mice by i.p. injection in 200µl PBS. 5-30 cysts from 4 week-infected CBA/J brain homogenate were then used to infect animals in all experiments.

### Immunohistochemistry

Mouse Tissue Immunofluorescence:

Reporter mice were perfused with 30 mL PBS followed by 30 mL 4% PFA (Electron Microscopy Sciences). All non-reporter strains were only perfused with PBS. Brains were cut along the midline and post-fixed in 4% PFA for 24h at 4°C. Brains were then cryoprotected in 30% sucrose (Sigma) for 24h at 4°C, embedded in OCT (Tissue Tek), and frozen on dry ice. Samples were then stored at −20°C. 40 µm sections were cut using a CM 1950 cryostat (Leica) and placed into a 24-well plate containing PBS. Sections were blocked in PBS containing 2% goat or donkey serum (Jackson ImmunoResearch), 0.1% triton, 0.05% Tween 20, and 1% BSA for 1h at RT. Sections were then incubated with primary antibody diluted in blocking buffer at 4°C overnight. Sections were washed the following day and incubated with secondary antibody in blocking buffer at room temperature for 1h. Sections were then washed and incubated with DAPI (Thermo Scientific) for 5 min at RT. Sections were then mounted onto Superfrost microscope slides (Fisherbrand) with aquamount (Lerner Laboratories) and coverslipped (Fisherbrand). Slides were stored at 4°C before use. Images were captured using an TCS SP8 confocal microscope (Leica) and analyzed using Imaris (Bitplane) software. Volumetric analysis was achieved using the surfaces feature of Imaris.

### Human tissue immunofluorescence

Slides containing 4 µm sections of human brain tissue were received from the UVA Biorepository and Tissue Research Facility and de-paraffinized in a gradient from 100% xylene (Fisher) to 50% ethanol (Decon Laboratories). Slides were then washed in running water and distilled water. Antigen retrieval was performed by incubating slides in antigen retrieval buffer (10 mM sodium citrate, 0.05%Tween-20, pH 6.0) in an Aroma digital rice cooker for 45 min at 95°C. Slides were then washed in running water followed by PBS-TW. Slides were then incubated with primary and secondary antibodies as described above for mouse brain tissue. Prior to imaging, Autoflourescence Eliminator Reagent was applied per the manufacturer’s instructions (EMD Millipore).

### Tissue processing and flow cytometry

Whole PBS-perfused brains were collected into 4 mL of complete RPMI (cRPMI)(10% fetal bovine serum, 1%NEAA, 1%Pen/Strep, 1%Sodium Pyruvate, 0.1%-β-mercaptoethanol). Papain digestion was performed for the chimera experiment. To perform papain digestion, brains were cut into 6 pieces and incubated in 5 mL HBSS containing 50U/mL DNase (Roche), and 4U/mL papain (Worthington-Biochem) for 45 min at 37°C. Tissue was triturated first with a large bore glass pipette tip, and twice with a small-bore pipette tip every 15 min. In all other experiments collagenase/dispase was used to digest brain tissue. To perform collagenase/dispase digestion, perfused brains were minced using a razor blade and passed through an 18-gauge needle. Brains were then digested with 0.227mg/mL collagenase/dispase and 50U/mL DNase(Roche) for 1h at 37°C. Following digestion, homogenate was strained through a 70 µm nylon filter (Corning). Samples were then pelleted and spun in 20 mL 40% Percoll at 650 g for 25 min. Myelin was aspirated and cell pellets were washed with cRPMI. Finally, samples were resuspended in cRPMI and cells were enumerated. Spleens were collected into 4 mL cRPMI and macerated through a 40 µM nylon filter (Corning). Samples were pelleted and resuspended in 2 mL RBC lysis buffer (0.16 M NH_4_Cl) Samples were then washed with cRPMI, and resuspended for counting and staining. In cases of acute infection, 4mL of peritoneal lavage fluid was pelleted and resuspended in 2mL of cRPMI for counting and staining. Single cell suspensions were pipetted into a 96 well plate and pelleted. Samples were resuspended in 50 μL Fc Block (0.1 µg/ml 2.4G2 Ab (BioXCell), 0.1% rat gamma globulin (Jackson Immunoresearch)) for 10 min. Cells were then surface stained in 50 μL FACS buffer (PBS, 0.2% BSA, and 2 mM EDTA) for 30 min at 4°C. Following surface staining, cells were fixed for at least 30 min at 4°C with a fixation/permeabilization kit (eBioscience) and permeabilized (eBioscience). Samples were then incubated with intracellular antibodies in permeabilization buffer for 30 min at 4°C. Samples were run on a Gallios flow cytometer (Beckman Coulter), and analyzed using Flowjo software, v. 10.

### qRT-PCR

Perfused brain tissue (100 mg) was placed into bead beating tubes (Sarstedt) containing 1mL Trizol reagent (Ambion) and zirconia/silica beads (Biospec). Tissue was homogenized for 30 seconds with a Mini-bead beater (Biospec) machine. RNA was extracted following homogenization per the Trizol Reagent manufacturer’s instructions. Complementary DNA was then synthesized using a High Capacity Reverse Transcription Kit (Applied Biosystems). Taqman gene expression assays were acquired from Applied Biosystems and are listed in the key resources table. A 2X Taq-based mastermix (Bioline) was used for all reactions and run on a CFX384 Real-Time System (Bio-Rad). Hprt was used as the brain housekeeping gene and relative expression to wildtype controls was calculated as 2^(-ΔΔCT)^.

### *T. gondii* cyst counts

Brain tissue (100 mg) was minced with a razor in 2mL cRPMI. Brain tissue was then passed through an 18 gauge and 22-gauge needle. 30 uL of resulting homogenate was pipetted onto a microscope slide (VWR) and counted on a Brightfield DM 2000 LED microscope (Leica). Cyst counts were extrapolated for whole brains.

### Bone marrow chimera

Wildtype B6.SJL-Ptprc^a^ Pepc^b^/BoyJ (C57BL/6 CD45.1) and ST2KO C57BL/6 mice were irradiated with 1000 rad. Irradiated mice received 3×10^6^ bone marrow cells from CD45.1 and CD45.2 donors the same day. Bone marrow was transferred by retro-orbital i.v. injection under isoflurane anesthetization. All mice received sulfa-antibiotic water for 2 weeks post-irradiation and were given 6 weeks for bone marrow to reconstitute. At 6 weeks, tail blood was collected from representative mice and assessed for reconstitution by flow cytometry. Mice were then infected for 4 weeks prior to analysis.

### Brain homogenate *ex vivo* supernatant collection

Brains were harvested from naïve and infected mice. Brain tissue was processed down to a single cell suspension as outlined in “tissue processing and flow cytometry”. Cells from half a brain were incubated in a 96-well plate in 200μL for 4 hours at 37°C. Cells were then pelleted at 1500rpm for 5 minutes, and supernatant was stored at −80°C for measurement of IL-33 by ELISA.

### CSF collection

Naïve and infected mice were anesthetized with ketamine/xylazine, their necks were shaved and obstructing skin and tissue surrounding the dura of the cisterna magna was removed. A small glass capillary was inserted into the dura of the cisterna magna, allowing approximately 10μL of CSF to fill the capillary. CSF from 4-5 mice was pooled and frozen at −80°C.

### IL-33 ELISA

The IL-33 Quantikine ELISA kit was purchased from Applied Biosystems and manufacturer instructions were followed. 50μL of either brain homogenate supernatant (*ex-vivo* assay) or CSF was used as sample volume. 4-5 CSF samples from individual animals were pooled to reach 50μL sample volume. IL-33 standards were serially diluted to a minimum of 30pg/mL. Final values of standards and samples were read on a spectrophotometer at 450nm.

### ACSA-2/CD11b magnetic enrichment

Brain tissue was harvested from mice and processed down to a single cell suspension as described in “tissue processing and flow cytometry” methods. The Anti-ACSA2 and CD11b Microbead kits, purchased from Miltenyi, were used to enrich for astrocytes and macrophages, respectively, over magnetic columns. In the case of enriching for astrocytes, both kits were used to first remove CD11b+ cells and then enrich for ACSA-2+ cells. Double-purifications were performed in all cases, which was optional, but recommended, by Miltenyi’s protocol. Cells were pelleted at 1500rpm and stored in 300μL Trizol (Ambion) at −80°C until further use.

### Statistical analyses

Statistical analyses comparing two groups at one time point were done using a student’s t-test in Prism software, v. 7.0a. Statistical analyses comparing more than two groups within the same timepoint or infection were done using a one-way Anova. In instances where data from multiple infections were combined, all from the same time-point post infection, a randomized block ANOVA was performed using R v. 3.4.4 statistical software to account for variability between infections. Genotype was modeled as a fixed effect and experimental day as a random effect. P values are indicated as follows: ns=not significant p>.05, * p < .05, ** p < .01, *** p < .001. The number of mice per group, test used, and p values are denoted in each figure legend. Data was graphed using Prism software, v.7.0a.

## FIGURES

**Figure S1.**
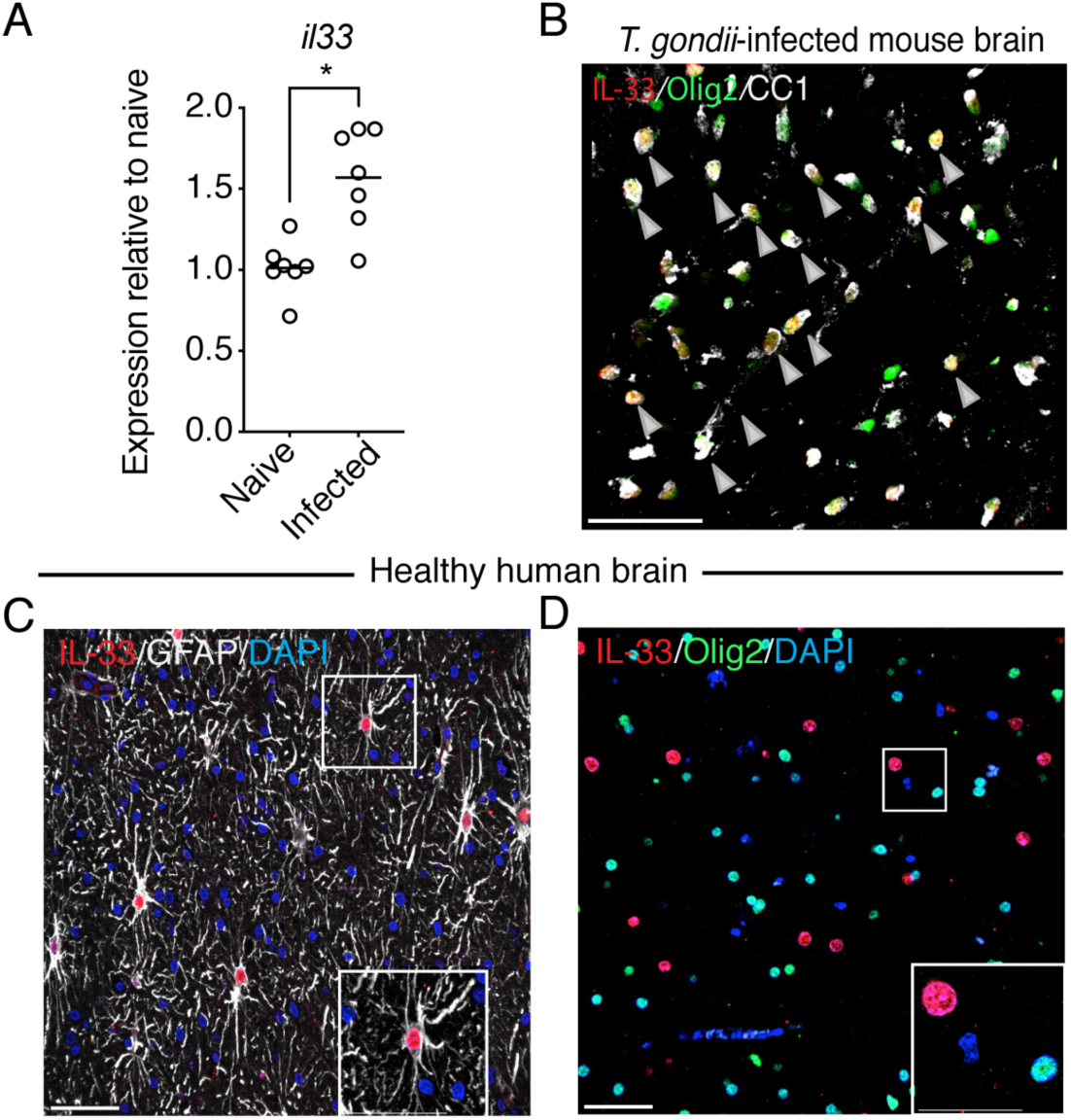
IL-33 expression in mouse and human brain tissue. **(A)** Real time PCR for IL-33 transcript from whole brain homogente at 4 weeks post infection compared to naïve brain tissue. **(B)** Colocalization, denoted by gray arrows, of nuclear IL-33 (red) with mature oligodendrocytes, marked by nuclear Olig2 expression (green), and CC1(white) by confocal fluorescence microscopy of infected mouse brain tissue. **(C and D)** Confocal fluorescence microscopy of nuclear IL-33 stain present in astrocytes (C) but not oligodendrocytes (D) in the healthy human temporal lobe. Statistical significance was determined by randomized block ANOVA (A), which shows data pooled from two independent experiments *= p<.05, **= p<.01, ***= p<.001. Scale bars indicate 50μm.

**Figure S2.**
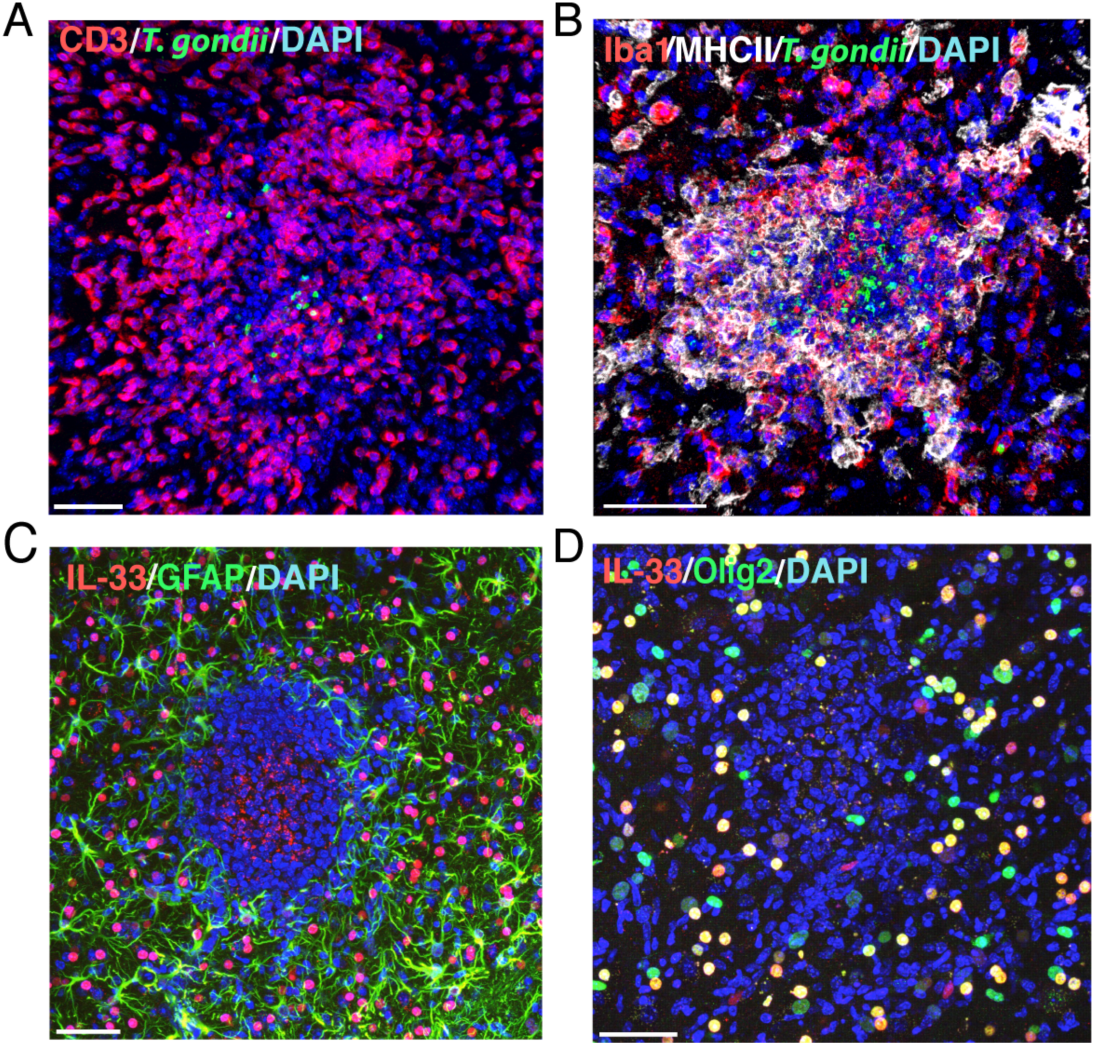
Inflammatory lesions in *T.gondii*-infected brain tissue. **(A and B)** Representative images of immune cells surrounding foci of individual replicating parasites (green) in cortical brain tissue, including CD3+ T cells (red) (A), and MHCII+ (white) Iba1+ (red) myeloid cells (B). **(C and D)** Representative images of necrotic foci, featuring a loss of brain resident cells which express IL-33 (red), including GFAP+ astrocytes (green) (C), and Olig2+ oligodendrocytes (green) (D). Scale bars indicate 50μm.

**Figure S3.**
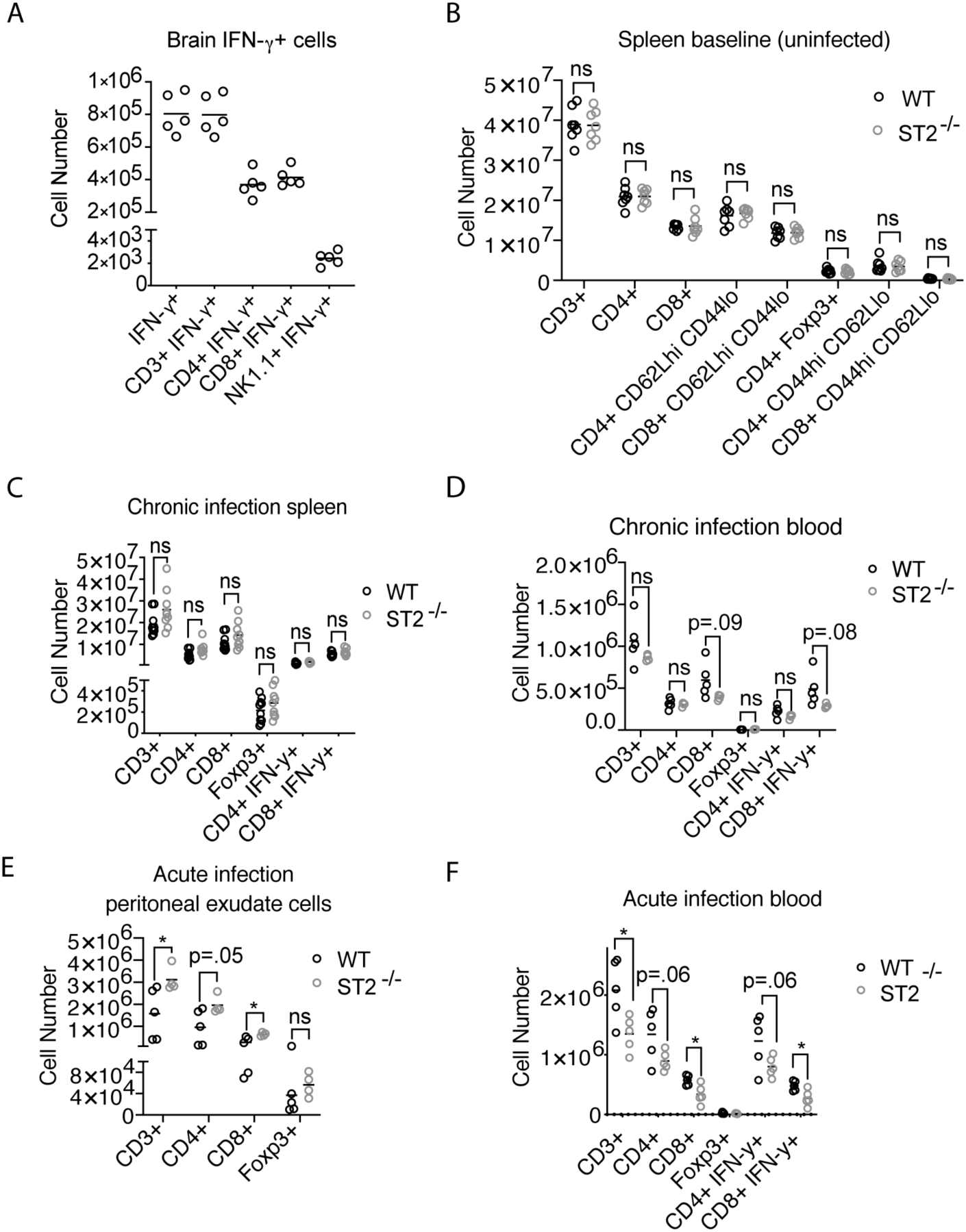
T cells in peripheral tissues and during acute infection in the absence of IL-33-ST2 signaling. **(A)** Breakdown of total IFN-γ+ cells by cell type by flow cytometry in infected brain tissue four weeks post infection. IFN-γ was measured following stimulation *ex vivo* for five hours with PMA/ionomycin. **(B)** Assessment of spleen T cell numbers in ST2-deficient mice prior to infection by flow cytometry. **(C and D)** assessment of peripheral tissue T cell numbers and activation, including spleen (C) and blood (D) by flow cytometry 4 weeks post infection. **(E and F)** T cell numbers at day 10 acute infection by flow cytometry in the peritoneum (E) and blood (F). Statistical significance was determined by randomized block ANOVA when two experiments were pooled (B and C), or by two-tailed t-test (D, E, F) *= p<.05, **= p<.01, ***= p<.001.

**Figure S4.**
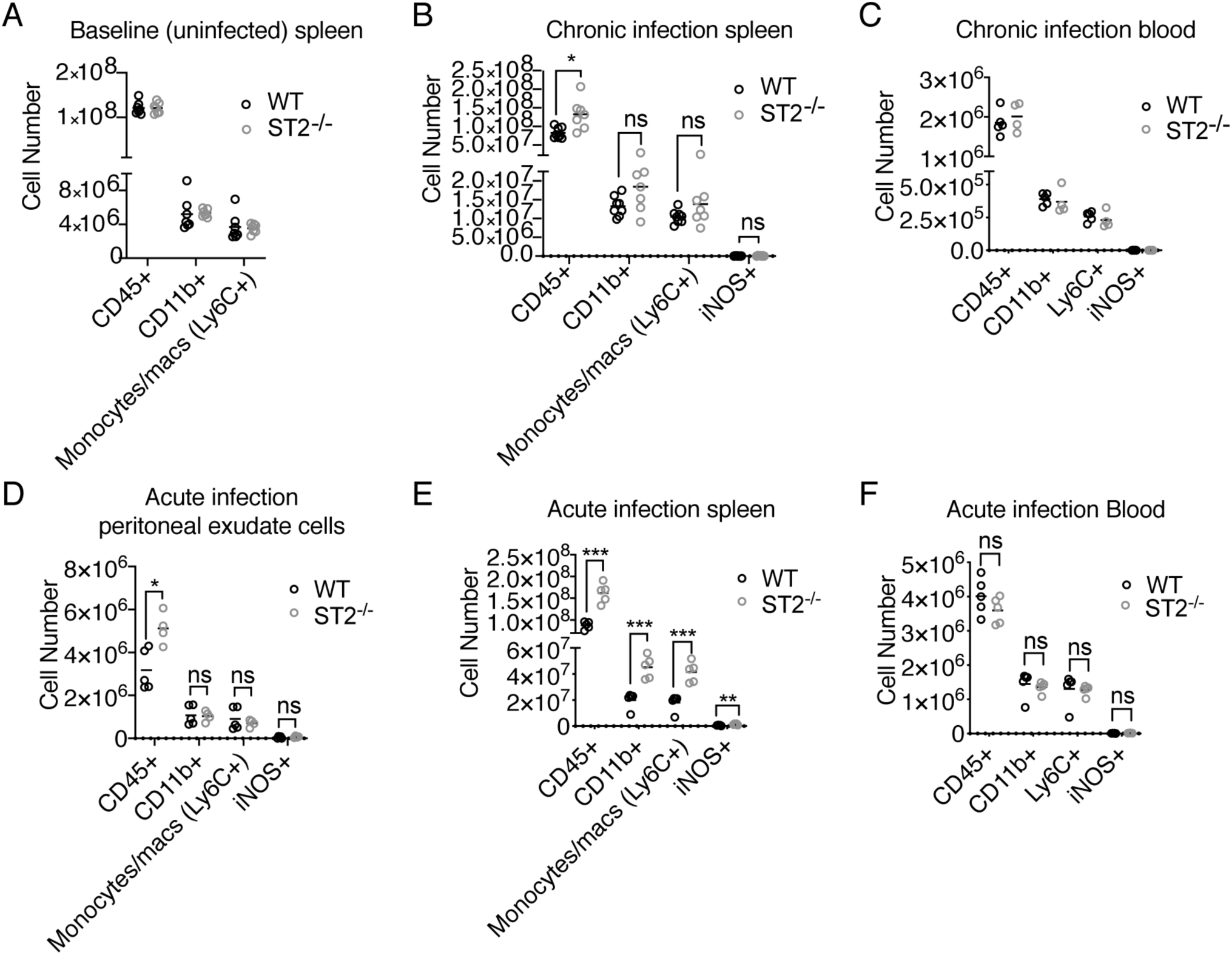
The myeloid cell response is intact in peripheral tissues and during acute infection in the absence of IL-33-ST2 signaling. **(A)** Assessment of spleen myeloid cell numbers in ST2-deficient mice prior to infection by flow cytometry. **(B** and **C)** assessment of peripheral tissue myeloid cell numbers and activation, including spleen (B) and blood (C) by flow cytometry 4 weeks post infection. **(D-F)** Myeloid numbers at day 10 during acute infection by flow cytometry in the peritoneum (D), spleen (E) and blood (F). Statistical significance was determined by randomized block ANOVA when two experiments were pooled (A and B), or by two-tailed t-test (C-F) *= p<.05, **= p<.01, ***= p<.001.

**Figure S5.**
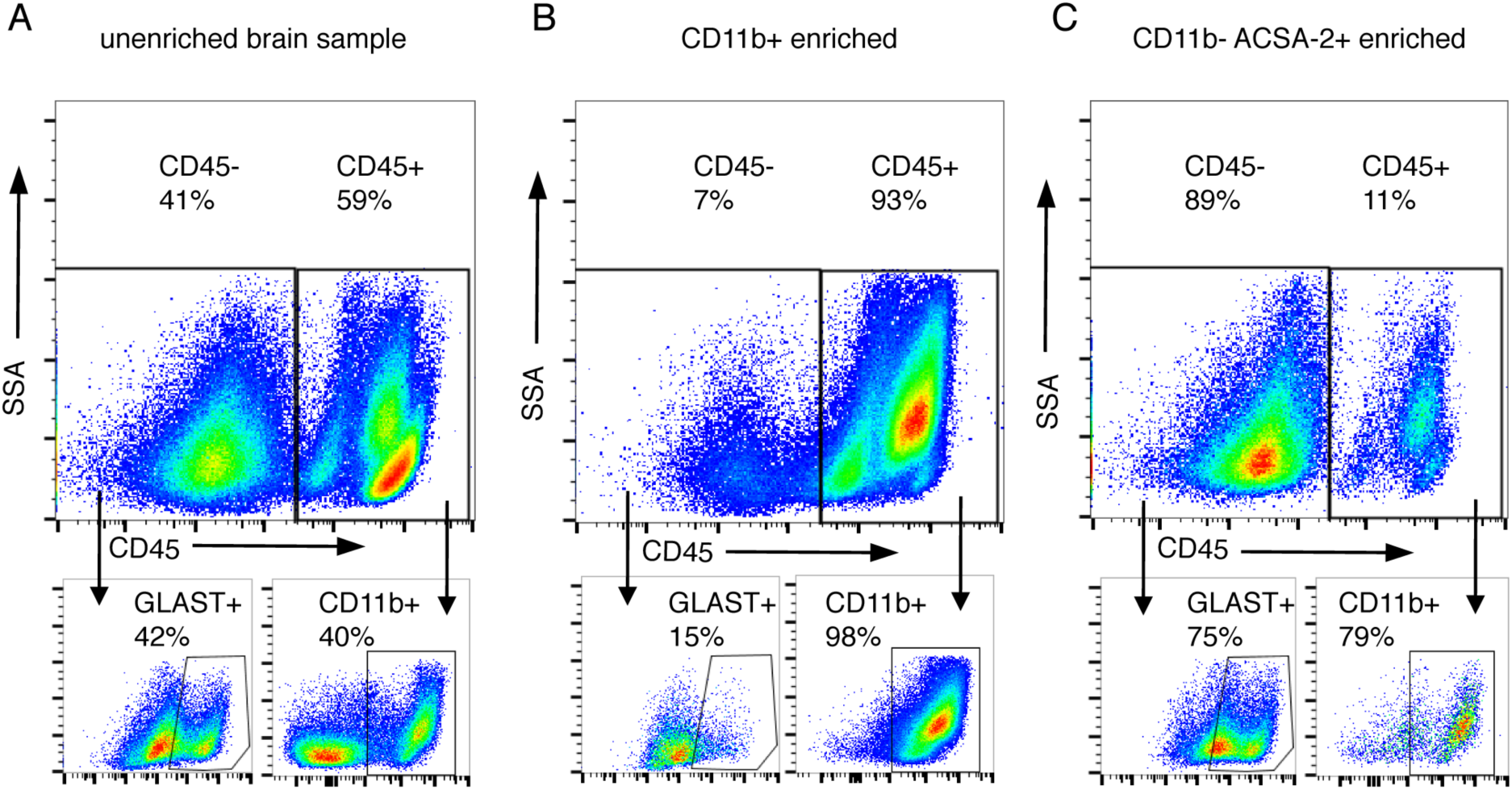
Magnetic enrichment for myeloid cells or astrocytes from infected brains. **(A)** Unenriched single cell suspension of all purified cells from infected brain tissue 4 weeks post infection. **(B and C)** Assessment of purity achieved by enriching for myeloid cells using CD11b+ magnetic beads (B), or astrocytes (C), by negatively selecting for myeloid cells using CD11b+ magnetic beads, followed by positive selection for astrocytes with ACSA-2+ magnetic beads.

**Figure S6.**
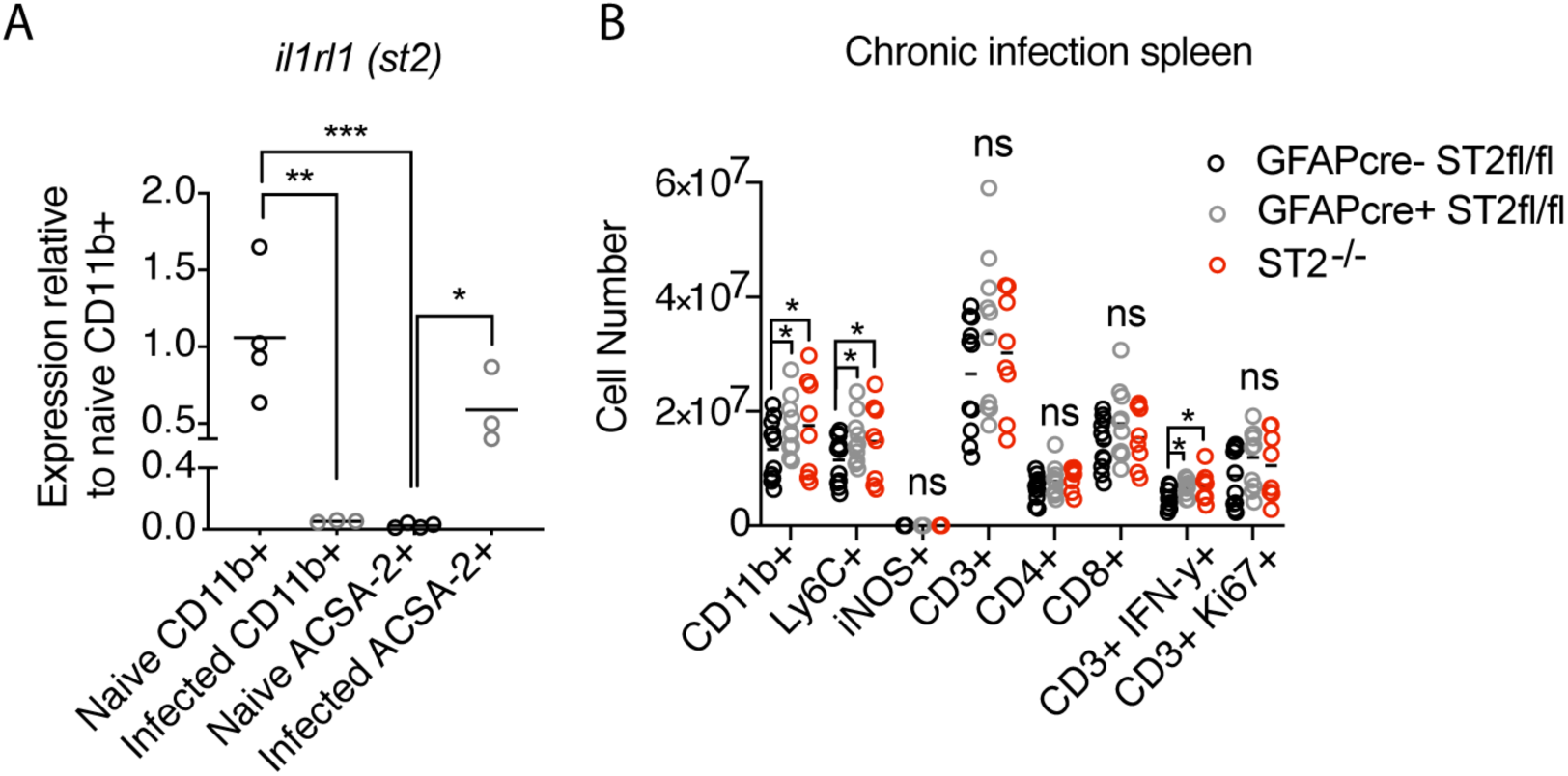
Astrocyte ST2 expression with infection, and astrocytic ST2-dependence for peripheral immune cell numbers. **(A)** Cell-type specific magnetic enrichment for myeloid cells (CD11b+) or astrocytes (CD11b- and ACSA-2+) in naïve and chronically infected brain tissue. Single cell suspensions of enriched cells were resuspended in Trizol, RNA extracted, and measured by real time PCR for *ilrl1*(st2) expression. **(B)** Assessment of spleen immune cell numbers by flow cytometry four weeks post infection. Statistical significance was determined by one-way ANOVA with Tukey’s test (A), or randomized block ANOVA (B) *= p<.05, **= p<.01, ***= p<.001.

## Notes

### Competing Interest Statement

The authors have declared no competing interest.

